# Long runs of homozygosity are correlated with marriage preferences across global population samples

**DOI:** 10.1101/2021.03.04.433907

**Authors:** Samali Anova Sahoo, Arslan A. Zaidi, Santosh Anagol, Iain Mathieson

**Author notes:** Correspondence to AZ or IM.

## Abstract

Children of consanguineous unions carry long runs of homozygosity (ROH) in their genomes, due to their parents’ recent shared ancestry. This increases the burden of recessive disease in populations with high levels of consanguinity and has been heavily studied in some groups. However, there has been little investigation of the broader effect of consanguinity on patterns of genetic variation on a global scale. Here, we collect published genetic data and information about marriage practices from 396 worldwide populations and show that preference for cousin marriage has a detectable effect on the distribution of long ROH in these samples, increasing the expected number of ROH longer than 10Mb by a factor of 1.5 (P=2.3 × 10^−4^). Variation in marriage practice and consequent rates of consanguinity is therefore an important aspect of demographic history for the purposes of modeling human genetic variation. However, marriage practices explain a relatively small proportion of the variation in ROH distribution and consequently the ability to predict marriage practices from population genetic samples (for example of ancient populations) is limited.

## Introduction

Marriage practices have consequences for human genetic variation. One extensively debated and regulated practice is consanguinity - the union between closely related individuals. An estimated 10.4 percent of the world’s population is made up of couples who are second-cousins or closer and their offspring [1] (see [2] and consang.net for additional estimates of worldwide consanguinity). The children of consanguineous marriages have longer runs of homozygosity [3] and therefore carry a greater recessive disease burden [4] equivalent to an increase in child mortality or severe disease of approximately 3-4% [5–7]. It is, however, unclear to what extent ethnographic assessments (either via surveys or direct observation) of marriage practices at the population level translate to the genetic signature associated with cousin marriage.

Although actual levels of consanguinity must be correlated with patterns of genetic variation, there are three reasons why ethnographic assessments of cousin marriage prevalence might not. First, different cultures’ understanding of “cousin” marriage could cover different concepts. For example, some might have a clear distinction between first and second cousin marriage, while others consider all of these types the same. Second, cultural preferences for marriage may change rapidly. Ethnographic data may be outdated, or preferences may not persist long enough to to have a detectable effect on patterns of genetic variation diagnostic of ongoing consanguinity. Finally, genomic signatures of consanguinity may be obscured by other demographic factors such as endogamy.

For these reasons, populations that practice cousin marriage may in fact demonstrate relatively few of the negative genetic consequences associated with increased homozygosity. For example, many marriages might occur between “cousins” who are actually quite distantly related. Alternatively, in many regions of the world, cousin marriage co-occurs with endogamy (marriage within defined sub-groups such as castes) [8]. Since endogamy can itself lead to excess recessive disease burden [9], the marginal effects of consanguinity might be small. Finally, even though both endogamy and consanguinity, by increasing ROH, increase the risk of recessive disease in the short term, the same process exposes such alleles to selection, purging them more effectively from the population in the long term.

Most previous work on the relationship between consanguinity and genetic variation has focused on a limited number of populations. McQuillan et al. 2008 correlated pedigree based measures of marriage practices to ROH in the Orcadian population [10]. Kang et. al. (2016) found a correlation between long ROH in recent individuals and inbreeding coefficients for nine Jewish populations, where rates of consanguinity were based on survey data [11]. Arciero et al. (2020) found a strong correlation between survey measured consanguinity and long ROH in a British Pakistani cohort [12]. In the most similar study to ours [13], Pemberton et al. (2014) found a correlation of .349 between inbreeding coefficients estimated from genetic data and survey-based consanguinity data. However, this analysis was based on only 26 populations and did not correct for correlations among populations due to shared demographic history.

We therefore set out to test whether a cultural preference for cousin marriage is detectable in genetic data from a worldwide sample of 3,859 individuals from 396 populations. We categorize these populations into those prohibiting (43%), allowing (12%), and preferring (44%) cousin marriage using publicly available ethnographic sources. We find that ethnographic measures of preferences for cousin marriage are detectably related to the distribution of long ROH, increasing the expected number of ROH longer than 10Mb by 1.5× in populations preferring cousin marriages over those that prohibit it (P = 2.31 × 10^−4^), after controlling for ten principal components of genetic variation, shorter runs of homozygosity, and heterozygosity (collectively serving as proxies for demographic events such as bottlenecks, admixture and endogamy). This effect corresponds to an approximate increase of 0.25 (in proportion of unions that are cousins) in populations where ethnographic data indicated a preference for the practice. Further, we find that populations preferring consanguinity show greater long ROH when compared to geographically and genetically “close” consanguinity-prohibiting populations in a matched-pair design.

## Methods

### Simulations

First, we carried out simulations to investigate the effect of degree (proportion of cousin unions) and duration (number of generations) of consanguinity on the distribution of runs of homozygosity. We also used these simulations to calibrate the difference in consanguinity between populations that we classified as prohibiting and preferring cousin unions (see below). We used cousin-sim [14] to generate pedigrees, each of which was initialized with 500 unrelated founders and 500 individuals in subsequent generations for either 10 or 50 generations. In each generation, (*c* ∈ {0,0.25,0.5,0.75}) of all unions were between cousins, of which 62% were first-, 23% were second-, and 15% were third-degree cousins [15]. We used the pedigree as input to *ped-sim* [16] with a Poisson crossover model and a refined genetic map of the human genome from [17] to simulate chromosomal segments and retained only those that were identical by descent (IBD) within an individual. We refer to these segments as runs of homozygosity (ROH) to distinguish them from inter-individual IBD.

We also calculated the expected number and total length of long runs of homozygosity analytically using the model described in [18]. Briefly, there are *m = 2n* **+ 4** meioses separating the two chromosomes of an offspring of *n^th^* cousins. For such an offspring, the number of segments of length *x* in the *i^th^* chromosome of length *l_i_* (in Morgans) has the following density (Equation S8 of [18]):

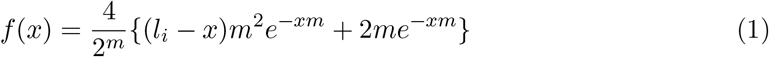

The integrals 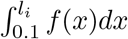 and 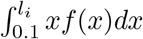 yield the expected number and total length, respectively, of long (>10cM or 0.1M) runs of homozygosity in the *i^th^* chromosome. This genetic length corresponds to approximately 10Mb physical distance, on average across the genome. We then computed the expected autosome-wide number and total length as 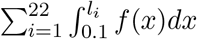 and 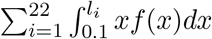, respectively, using the sex-averaged genetic map from [17] for chromosomal lengths (in Morgans).

### Genetic data collection and processing

We collected genotype data generated with the Affymetrix Human Origins array from 6 papers [9, 19–23], and merged this with whole-genome sequence data from three further papers [24–26]. We merged all datasets and removed 1) apparent duplicate samples (identified using *plink╌genome*) 2) samples marked as questionable in the original publications 3) samples where the population label was too generic to be useful. We retained the population labels from the publications except where there was an obvious typo or synonymous population. Our genetic dataset contained 4544 individuals from 488 populations (median number of individuals per-population 7, range 1-71). We restricted to autosomes, leaving 469,421 SNPs.

For each individual, we calculated heterozygosity (at genotyped SNPs) and the number of ROH greater than 1,2,5 and 10 Mb (using the *╌het* and *╌homozy* commands in *plink*). We then subtracted the number of ROH so that our variable NROH1 represents the number of ROH longer than 1 but smaller than 2 Mb, NROH5 representing the number of ROH longer than 5 but smaller than 10 Mb, and so on. Finally, we computed principal components of the merged dataset, and also separately of individuals belonging to the 237 South Asian populations. We used *plink* v1.90 [27] and *smartpca* v16000 [28] for all analyses.

### Ethnographic data collection and processing

We collected data on marriage practices from the following sources:

- The electronic Human Relations Area Files (eHRAF) World Cultures database [29], which catalogs a large set of ethnographic writing describing cultural and social aspects of different groups.
- The Ethnographic Atlas [30], which describes the cultural practices of 1,291 societies.
- Focused online searches for marital practices in specific groups. Sources include Google Scholar, Google books, as well as other papers and books. To search for specific populations, we searched for terms such as ‘marriage’, ‘cousin marriage’, ‘endogamy’, ‘exogamy’, ‘consanguinity’, and ‘consanguineous’ along with the name of the population.

We encoded the level of consanguinity in a population categorically as ‘prohibited’ (coded as 0), ‘permitted’ (1), or ‘preferred’ (2). When quantitative measures (e.g. prevalence of cousin unions in the population) were available, we designated populations to the above three categories based on the percent of unions between first cousins and first cousins once removed. In such cases, a prevalence of 0 to 2.5% was encoded as prohibited, 2.5% to 20% was encoded as ‘permitted’, and 20% and above was encoded as ‘preferred’. When quantitative measures were not available, we analyzed ethnographic records for terms indicating the preference for cousin unions. Groups that stated cousin marriage was “common”, “practiced”, “encouraged” or equivalent were classified as ‘preferred’ category. Meanwhile, groups where cousin marriage was described as being “allowed”, “occasional”, “present but uncommon” or equivalent were coded as ‘permitted’. Lastly, groups where cousin unions were “forbidden”, “barred” or marriage was “exogamous” were classified as ‘prohibited’. We classified groups where information was unclear or unavailable as ‘missing’. We gathered marriage practice information for 522 populations, which was reduced to 396 (and 3,859 individuals) after merging with the genetic data.

To validate our consanguinity scoring, we tested whether our assignments were correlated with individual survey data on marriage practices from the 2011 India Health and Development Survey (IHDS) [31]. We matched populations of individuals from the IHDS survey to population names in our genetics data using caste names provided in the IHDS data (N = 198 for the matched sample of Indian populations). Our consanguinity assignments are strongly correlated with the proportion of IHDS respondents answering yes to the following prompt: “Now, I would like to ask you some questions about marriage customs in your community (jati) for a family like yours. Is it permissible to marry a girl to her cousin?” (Fig. 1).

**Figure 1:**
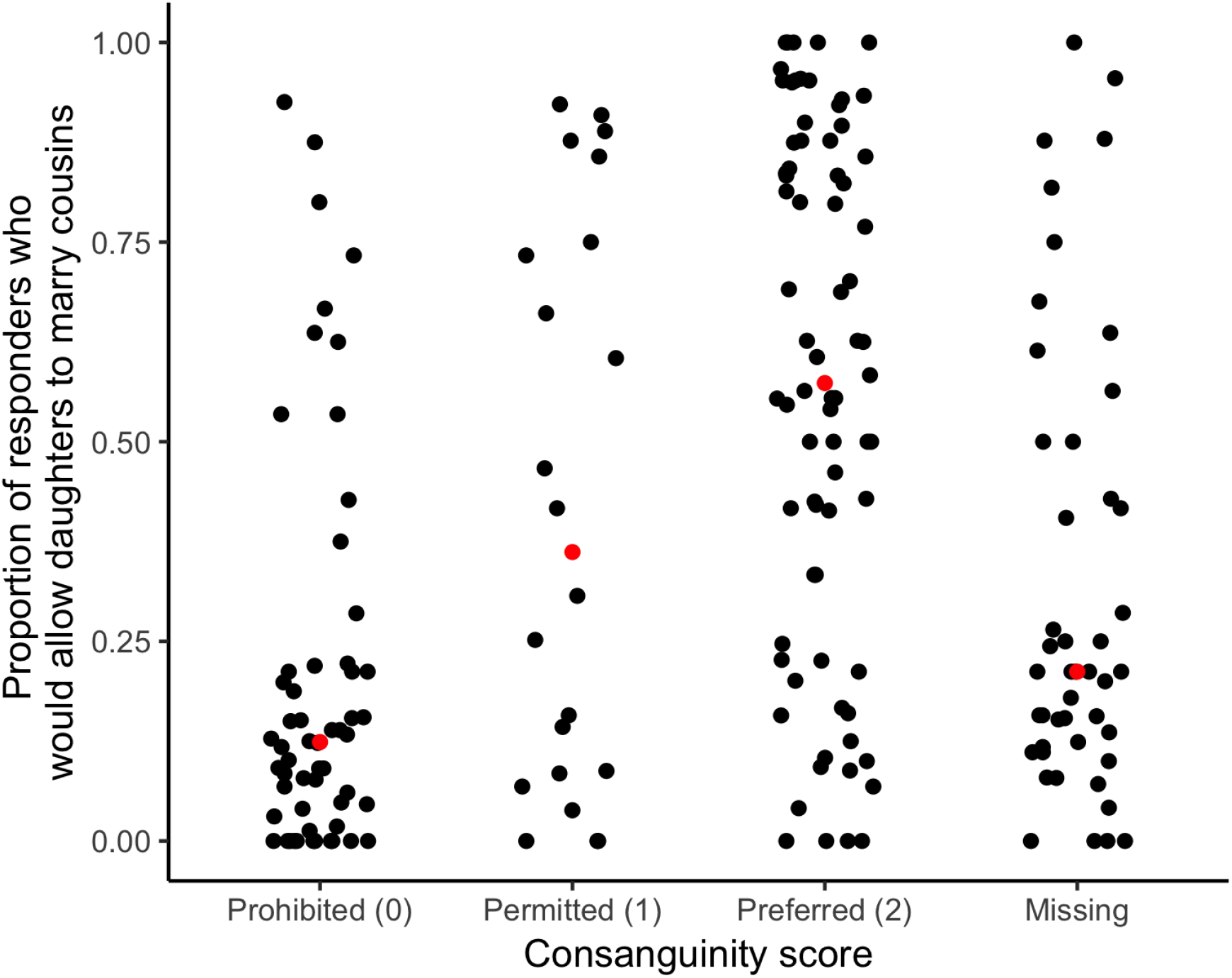
Our consanguinity score corresponds well with self-reports on whether cousin marriage is permitted in an individual’s community. The x-axis shows the consanguinity score assigned to each population and the y-axis shows the proportion of survey participants in the 2011 Indian Health and Development survey (IHDS) who responded that in their community it was permissible for a girl to marry their cousin. Each point is a population (matched using the caste name variable from the IHDS survey, N = 198) and red points represent the median for that category.

### Effect of consanguinity on long runs of homozygosity

We used Poisson regression to model the effect of consanguinity on the number of long (>10cM) runs of homozygosity (NROH). The consanguinity score of the population was treated as a categorical variable with three levels (‘prohibited’ - 0, ‘permitted’ - 1, and ‘preferred’ - 2). In all models, we included genome-wide heterozygosity, ten genetic principal components (gPC 1-10), and the number of shorter (1-2cM and 2-5cM) runs of homozygosity (hereafter NROH1 and NROH2, respectively) as covariates to correct for other demographic events such as population bottlenecks.

We fitted models where each population was an observational unit as well as models where each individual was treated as an observational unit. When each population was treated as an observational unit, we used the mean value of the variable (e.g. NROH and gPCs) in the population and fitted both weighted (weights proportional to the sample size from each population) and unweighted models. When each individual was treated as the observational unit, we used individuals’ values of NROH and gPC and population-wide consanguinity scores. Finally, because our densest sampling is from South Asia, we also fitted models restricting to populations from this region.

We evaluated model fit based on the dispersion parameter and the distribution of standardized residuals. In the main text, we present results from the Poisson model, which assumes that the mean of the residuals is equal to the variance. Indeed, the residuals showed no evidence of over-dispersion (Pearson statistic = 259.62, p-value under χ^2^ with 378 degrees of freedom ≈ 1).

### Paired analysis

To further ensure that the relationship between consanguinity and NROH10 was not confounded by long-term demographic history, we identified population pairs that were matched genetically and geographically but were different in that cousin unions were preferred in one population and prohibited in the other. To maximize genetic similarity and geographic proximity in selecting population pairs, we calculated genetic (Euclidean) distance using PC1-10 and geographic distances (using longitude and latitude) between populations that prefer consanguinity (score = 2) and those that either prohibit it or permit it (score ∈ {**0**,**1**}). Then, we divided each distance matrix by its median (to account for differences in scale) and averaged the two to calculate a single distance matrix. We selected population pairs in ascending order of the distance between them such that each population was part of only one pair. We used a maximum distance cutoff between populations (determined visually) to ensure that remaining populations did not pair with populations that were genetically and geographically distant. We used a Wilcoxon signed-rank test to test for a mean difference in NROH10 between population pairs.

### Predictive model

To predict consanguinity score from genetic data, we fitted a logistic regression model where consanguinity score (2 versus 0 or 1) was treated as the response and NROH10, NROH5, NROH2, NROH1, and heterozyosity were used as predictor variables. The performance of the predictive model was evaluated using a leave-one-out approach.

## Results

### Expected effect of consanguinity on ROH

Based on the analytical result derived in [18], we expect, on average, ≈ 6 ROH longer than 10cM in a population where all unions were between cousins; ≈ 9 if all of them are first cousins. Our simulations confirm that both the average total length and the number of ROH (of all lengths) increase as a function of increasing degree of consanguinity as well as the number of generations for which the practice continues. The simulation results also match well with the theoretical expectation derived for a single generation of cousin mating (black points in Fig. 2). In fact, the number of generations (either 1, 10, or 50) has little impact on the average number or total length of long (>10 cM) ROH (ROH10) (Fig. 2). This is because long runs of homozygosity arise primarily due to cousin unions in the recent past as long segments of the genome do not survive for many generations because of recombination. Therefore, long runs of homozygosity are only informative about consanguinity occurring in the very recent past. In contrast, the number of generations has a much larger impact on the number and length of all ROH, which are sensitive to long-term demographic history [3]. The degree of consanguinity (% unions that are between cousins) affects both long and short runs of homozygosity (Fig. 2).

**Figure 2:**
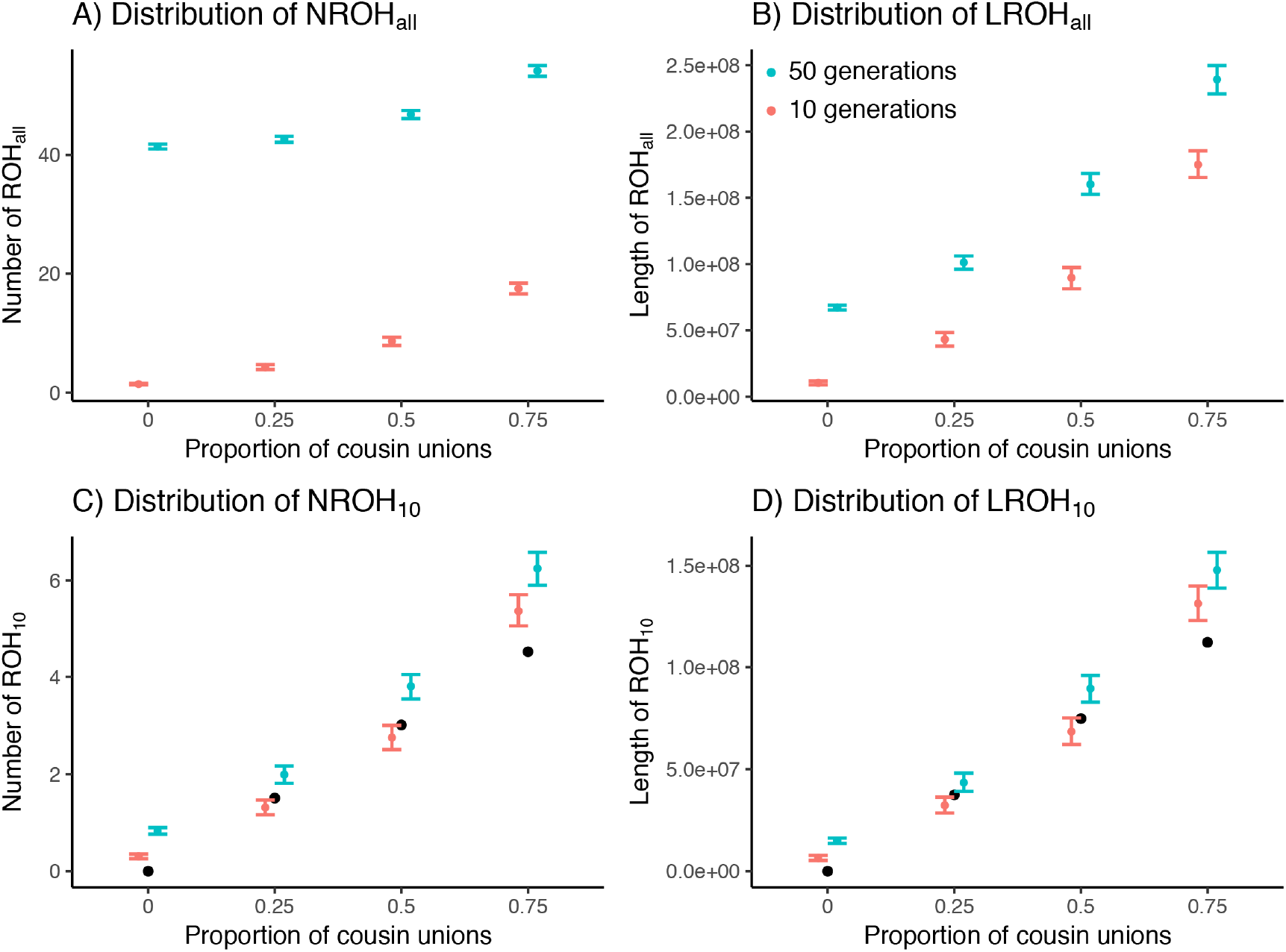
Distribution of the mean number (A and C) and length (B and D) of runs of homozygosity observed in pedigrees simulated with varying degree of consanguinity (proportion of unions between cousins) and number of generations for which the practice persisted — either 10 (red) or 50 (blue). Colored points and whiskers represent the mean and 95% bootstrap confidence interval, respectively, for (A and B) all runs of homozygosity and (C and D) for long (>10cM) runs of homozygosity. Black points represent the theoretical expectation from one generation of consanguinity.

### Geographic variation in cultural preference of consanguinity

We show the geographic variation in cultural preference for consanguinity in Fig. 3. It is interesting to note that in India, preferences for consanguinity appear to markedly change when comparing northern populations to southern populations (Figure 3B). Consanguinity is more prevalent in southern India compared with northern India. More formally, using linear regression, where the consanguinity score was used as a continuous variable, we show that prevalence of consanguinity varies significantly (ANOVA p-value = 1.22 × 10 ^−6^) between broad geographic regions (e.g. East Asia, Northern Africa, and Western Europe). Multinomial ordinal regression, where consanguinity score was treated as an ordinal variable, showed a similar result (p-value = 1.58 × 10 ^−6^). Two regions showed statistically significantly different preferences for consanguinity compared to the average. The Middle East and North Africa predicted a greater preference toward consanguinity (mean consanguinity is 1.67 on our scale from 0 to 2) whereas Western Eurasian populations on average were expected to have a lower preference for consanguinity (mean consanguinity of 0.53).

**Figure 3:**
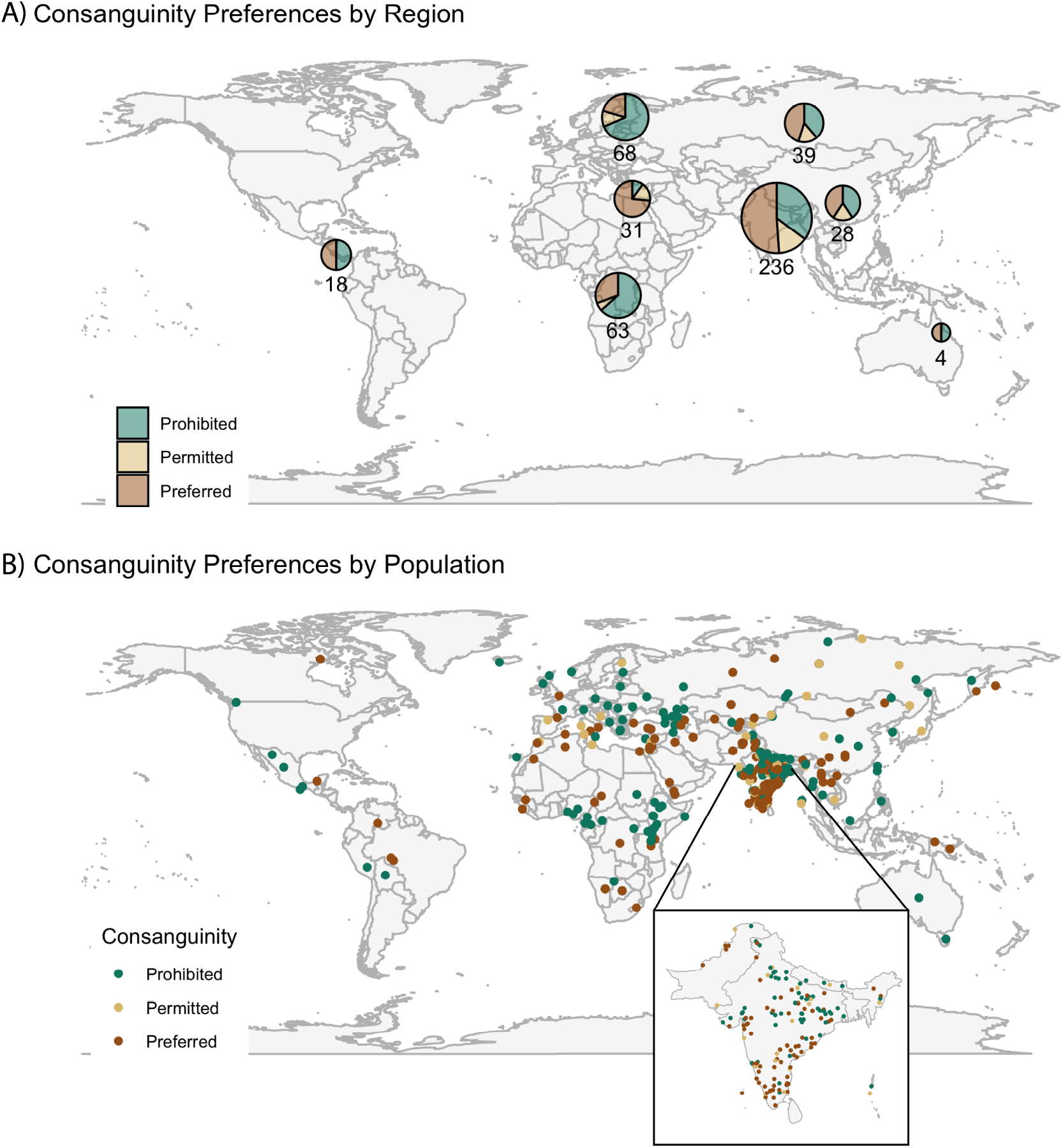
Graphical summary of prevalence of consanguinity across the world.

### Genetic footprint of cousin marriages

The number of long (>10cM) runs of homozygosity (NROH10) was positively associated with the degree of consanguinity when each individual was treated as an observational unit (N = 3,859 individuals, *β_score=2_* = 0.39, p-value = 3.02 × 10^−14^, Table 2, Fig. 4A). This coefficient can be interpreted as a 1.5 × increase in the number of long runs of homozygosity in populations with a strong preference for cousin marriage compared to populations where such unions are prohibited. The estimated effect is consistent if we treat populations (as opposed to individuals) as observational units (N = 396 populations, unweighted *β_score=2_* = 0.52, p-value = 2.31× 10^−4^; weighted *β_score=2_* = 0.60, p-value = 1.0× 10^−4^; Fig. 4B; Tables S1 and S2). It is also consistent when we restrict the analysis to populations within South Asia, a region that is very diverse in marriage practices and where we have the largest density of observations (individual model with N = 1363 individuals, *β_score=2_* = 0.59, p-value = 8.68 × 10^−10^; Tables S3, S4, S5; population model with N = 234 populations, unweighted *β_score=2_* = 0.49, p-value = 0.0415; weighted *β_score=2_* = 0.62, p-value = 0.034).

**Table 1:** Consanguinity prevalence by region (0 = prohibited, 1 = accepted, 2 = preferred).

|  | region2 | 0 | 1 | 2 | missing |
| --- | --- | --- | --- | --- | --- |
| 1 | Americas | 8 |  | 8 | 2 |
| 2 | Central Asia & Siberia | 12 | 5 | 14 | 8 |
| 3 | East Asia | 11 | 5 | 11 | 1 |
| 4 | Middle East & North Africa | 3 | 5 | 22 | 1 |
| 5 | Oceania | 2 |  | 2 |  |
| 6 | South Asia | 64 | 25 | 93 | 54 |
| 7 | Sub-Saharan Africa | 33 | 3 | 16 | 11 |
| 8 | Western Eurasia | 37 | 6 | 11 | 14 |

**Table 2:** Poisson regression model (individual) with NROH10 used as response. Coefficients for consanguinity score is shown relative to the score of 0 (consanguinity prohibited). Coefficients of 1 and 2 represent populations which permit and prefer cousin unions, respectively. Regional coefficients are shown relative to the Americas. Ten genetic PCs (calculated for the full sample) were also included in the model but their coefficients are not shown.

| | $\beta$ | SE($\beta$) | z value | Pr(>\|z\|) |
| --- | --- | --- | --- | --- |
| (Intercept) | 19.4099 | 0.4849 | 40.03 | $< 2 \times 10^{-16}$ |
| N.ROH1 | -0.0230 | 0.0020 | -11.76 | $< 2 \times 10^{-16}$ |
| N.ROH2 | -0.0105 | 0.0032 | -3.25 | 0.0012 |
| HET | -53.2682 | 0.9573 | -55.64 | $< 2 \times 10^{-16}$ |
| Consanguinity score = 1 | 0.1834 | 0.0625 | 2.93 | 0.0033 |
| Consanguinity score = 2 | 0.3945 | 0.0519 | 7.60 | $3.02 \times 10^{-14}$ |
| Region, Central Asia & Siberia | -0.0794 | 0.3244 | -0.24 | 0.8066 |
| Region, East Asia | -0.7236 | 0.3421 | -2.12 | 0.0344 |
| Region, Middle East & North Africa | -0.6708 | 0.3439 | -1.95 | 0.0511 |
| Region, Oceania | 0.3227 | 1.0099 | 0.32 | 0.7493 |
| Region, South Asia | -0.4938 | 0.3396 | -1.45 | 0.1459 |
| Region, Sub-Saharan Africa | 0.1513 | 0.5604 | 0.27 | 0.7872 |
| Region, Western Eurasia | -1.1983 | 0.3351 | -3.58 | 0.0003 |

**Figure 4:**
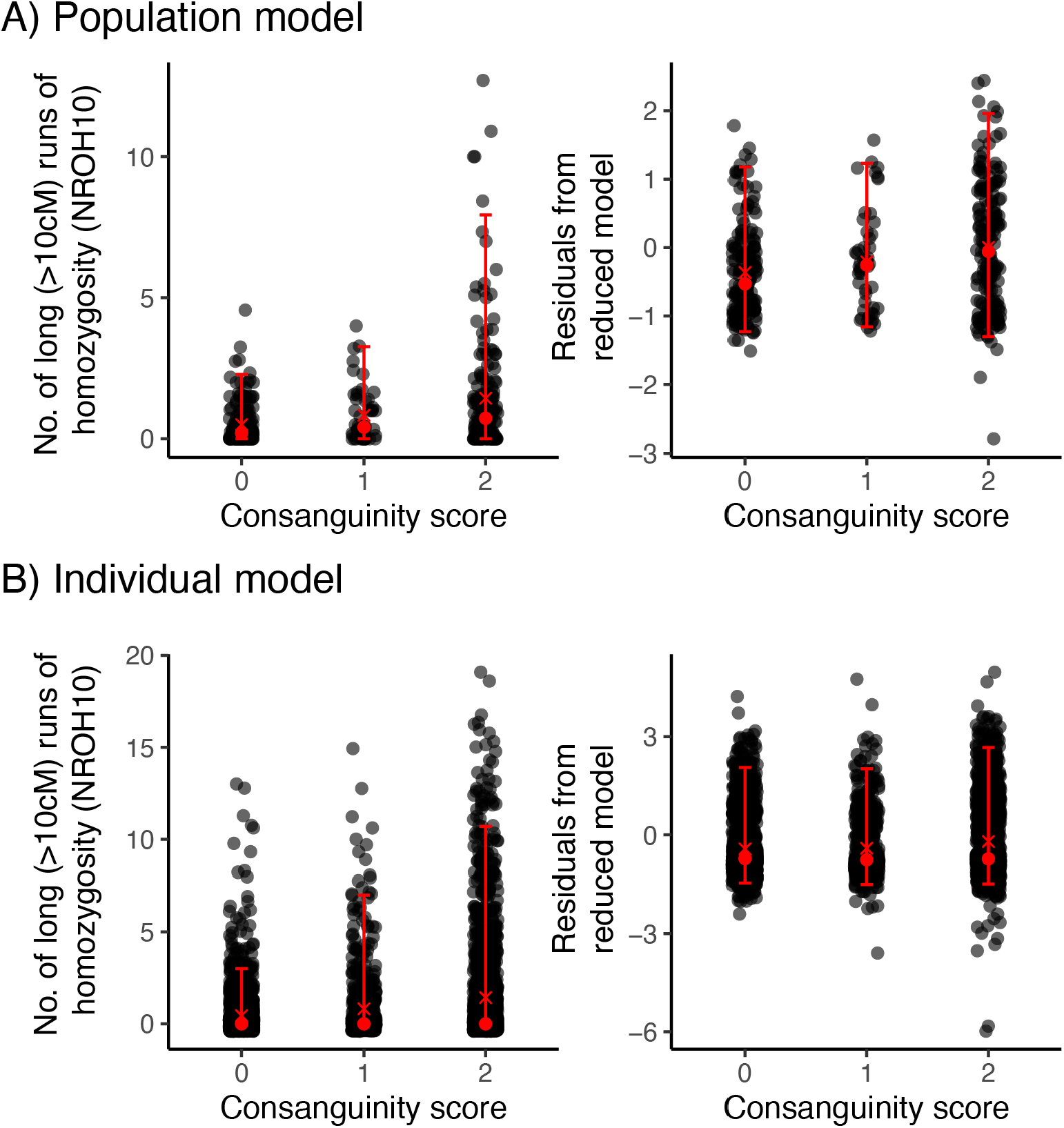
Relationship between consanguinity and NROH10. Each point is a population in the top row (A) and an individual in the bottom row (B). Panels in the first column show raw values of NROH10 on the y-axes and panels in the second column show the residuals of a reduced model where consanguinity score has been left out. Black points represent individual observations and red points represent median (point) and mean (cross) and horizontal bars represent the 95% CI.

Because the covariates in our regression models may not completely capture the effects and interactions of shared ancestry and culture, we confirmed this result with a matched-pair analysis. We identified pairs of populations matched for both genetic similarity and geographic proximity, but discordant for cousin marriage practice, and then tested for a difference in NROH within pairs (Methods). We find that NROH10 is significantly different between such matched populations (P = 0.007, N = 72 pairs) (Fig. 5) whereas other variables that are more sensitive to long-term demographic history such as NROH1, NROH2, NROH5, and HET are not (Table 3), consistent with results from the linear models. Removing populations with mean NROH10 > 5 did not change this result (Table S6).

**Figure 5:**
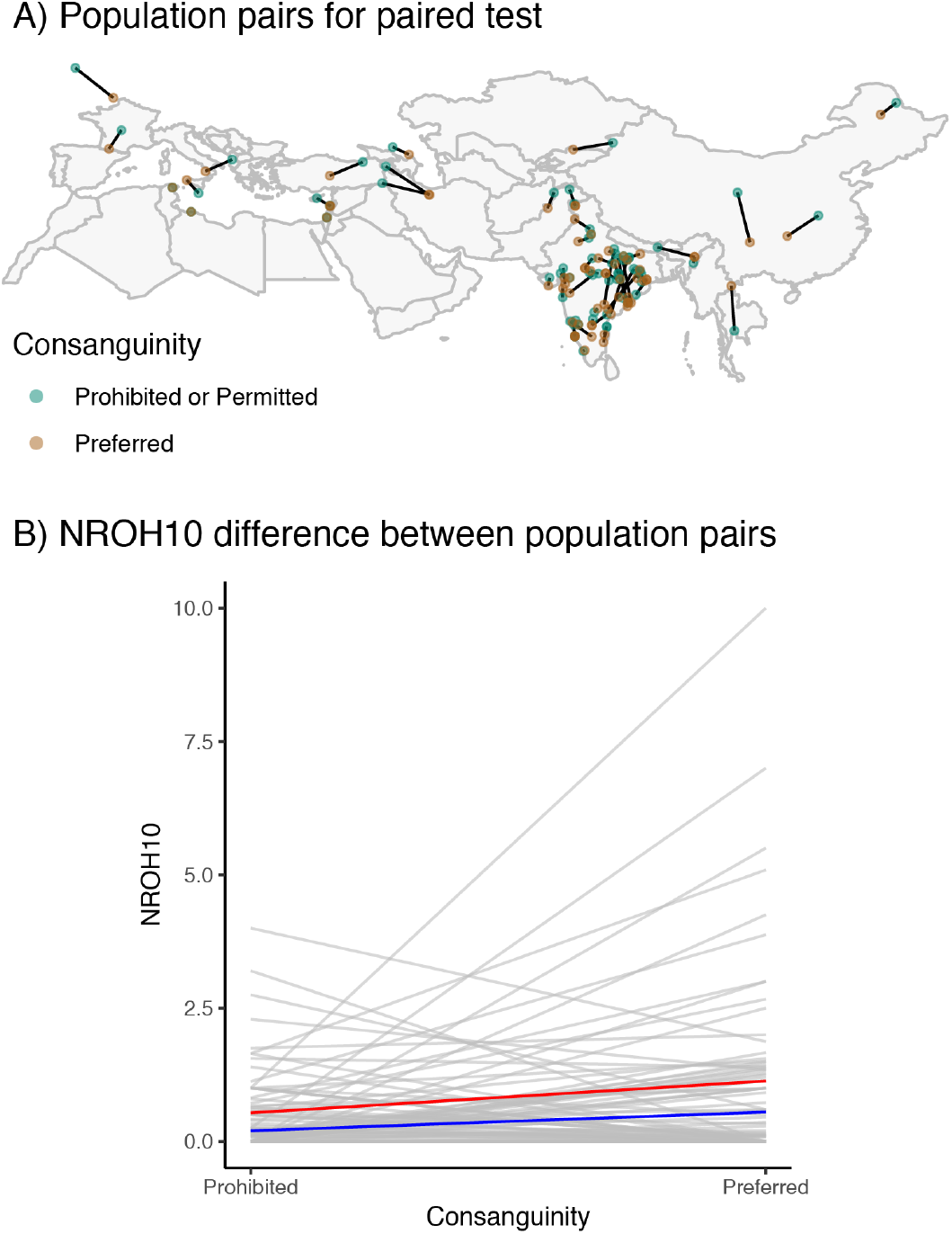
Paired comparison of NROH10 between populations where consanguinity is prevalent vs populations where it is not (N = 77 pairs). (A) Population pairs matched on genetic similarity and geographic similarity. Two pairs (one observed in Central America and one in North Africa) are not shown to maximize resolution in regions with a high density of pairs. (B) The mean number of long (>10cM) runs of homozygosity (NROH10) is greater in populations that prefer cousin unions compared to populations that prohibit them. The black lines demonstrate the comparison between pairs of populations that are geographically and genetically similar while the red and blue lines show the mean and median trend, respectively.

**Table 3:** Genetic differences between pairs of populations preferring and prohibiting consanguinity (N = 72 pairs)

|  | Variable | Statistic | Pvalue |
| --- | --- | --- | --- |
| 1 | NROH10 | 1405.50 | 0.007 |
| 2 | NROH5 | 1369.50 | 0.459 |
| 3 | NROH2 | 1280.00 | 0.851 |
| 4 | NROH1 | 1118.50 | 0.274 |
| 5 | HET | 1025.00 | 0.105 |

### Predicting marriage practice from genetic data

Finally, we asked to what extent it is possible to predict marriage practices based on the genomic distributions of ROH. We predict the binary variable of prohibited versus permitted or preferred cousin marriage using logistic regression using NROH1,2,5 and 10, and heterozygosity. We do not include region or principal components in the regression, which is conservative in this context. Using leave-one-out cross-validation we find that classification power is low (area under the curve, or AUC = 0.62) but well calibrated for predicted probabilities > **50%** (Fig. 6).

**Figure 6:**
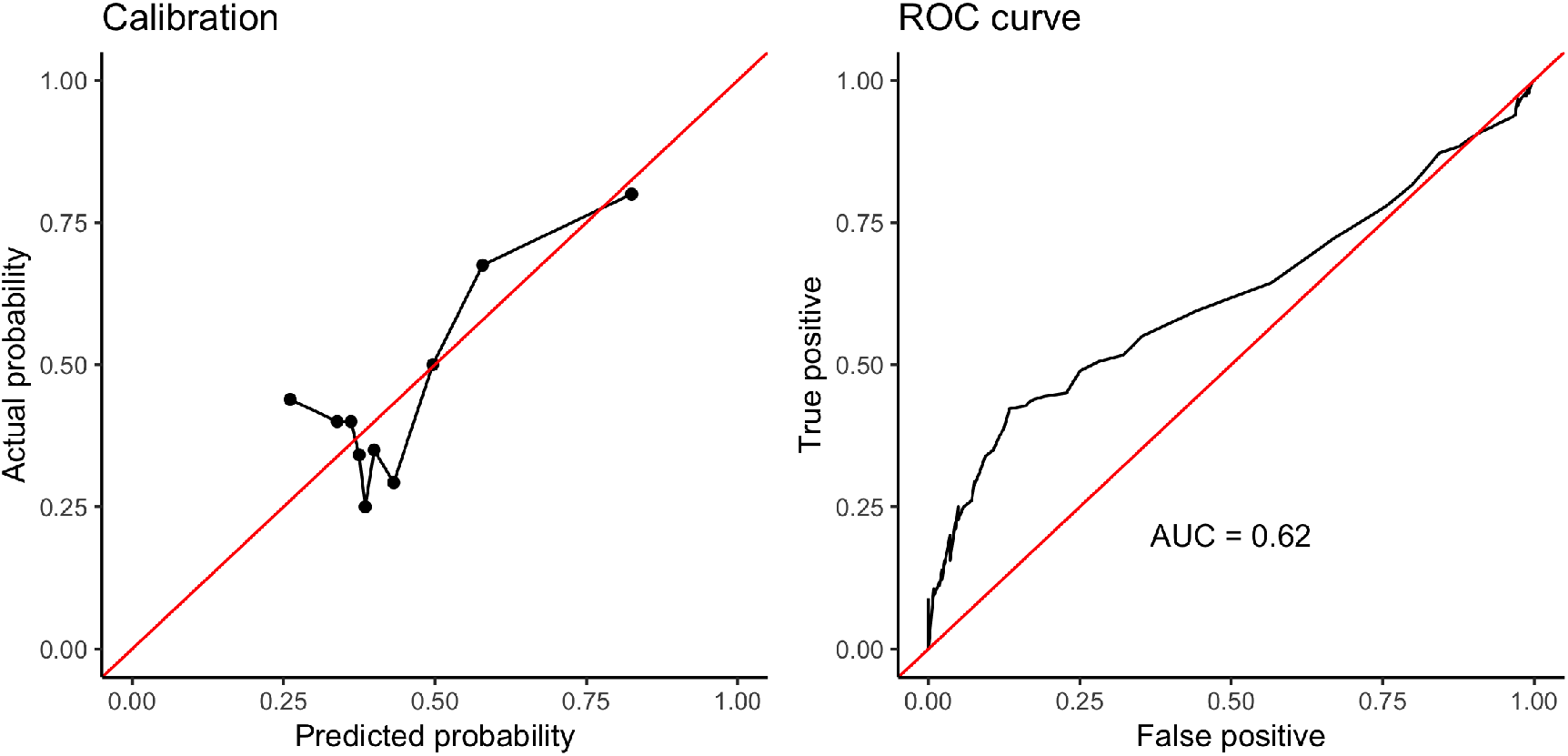
Results from predictive model. The model is well-calibrated (left) and the area under the ROC as the prediction cutoff varies is 0.62 (right).

## Discussion

The negative genetic consequences of marriage between close relatives have been well-known for centuries, yet the practice continues amongst millions of households around the world. Consanguinity creates long runs of homozygosity in the genome, increasing the risk of recessive disease in the offspring. We find a statistically significant relationship between ethnographic assessments of cousin marriage practice and long ROH in 3,859 individuals across 396 worldwide populations. Based on our simulations, this suggests that populations that prefer cousin marriage have, on average, actual first cousin marriage rates of approximately 25 percent (if we assume populations that prohibit cousin marriage have zero percent cousin marriages). This translates into an increase in child mortality or severe genetic disease of less than 1%—a cost that may in many cases be outweighed by the social and economic benefits of cousin marriage.

Indeed, social scientists have primarily focused on the benefits of cousin marriage, such as keeping land within the family line [32, 33], creating greater assurance that a marrying daughter will receive better treatment in her new household [7, 34], marrying a daughter to her cousin as barter for obtaining a cousin-bride for a son [35], or parents having a strong preference for the social status of their in-laws [36]. Less emphasis has been placed on integrating these benefits in to a broader cost-benefit framework that ultimately determines the emergence, persistence and potential decline of cousin marriage practice at the population level. Our results suggest that obtaining the benefits of cousin marriage does not appear to come with a large population level genetic cost. This is consistent with the long-standing prevalence of cousin marriage preference throughout the world, and suggests that future work incorporate genetic data, which might provide more accurate estimates of actual prevalence of consanguinity, as a cost-metric for this practice.

Our study has a number of limitations—some unavoidable. Our classification of marriage practices is based on literature search and many of our classifications may be incorrect or fail to reflect complexity or structure within named groups. Similarly, the genetic data we assembled typically have little information about sampling. Group labels may be misleading or incorrect and we may have linked them incorrectly to the ethnographic data. In addition since we are, to first order, measuring whether the individuals in our sample have consanguineous parents, our results could be biased if the sampled individuals were not representative of the wider populations. With more detailed data, many of these limitations can be avoided (e.g. [11]), although at the cost of limiting the geographic and cultural scope of the study. Despite these limitations, we do detect the expected associations, indicating that we are capturing an important part of the effect, although qualitative estimates of magnitude could be underestimated.

Finally, we also briefly explored the potential of using the present-day relationship between long ROH and population level assessments of cousin marriage to predict the marriage practices of ancient populations using ancient DNA. This has the potential to track the prevalence of consanguinity through time, especially for populations with sparse archaeological and historical records. Our results show that while high levels of ROH predict cousin marriage well, intermediate and low levels had little predictive power. This suggests that it may be difficult to predict population level preferences for cousin marriage using small samples of ancient DNA. Larger samples, more accurate predictors and integration of archaeological data might improve this prospect. In general, this kind of broad-scale survey of genetic data can be integrated with other types of information to address further questions around the existence, history, and the health and economic effects of consanguinity.

## Acknowledgements

This work was supported by NIGMS award number R35GM133708. The content is solely the responsibility of the authors and does not necessarily represent the official views of the National Institutes of Health.

---

**Table S1:** Poisson regression model fitted for NROH10 as response (population - unweighted). Ten genetic PCs (calculated for the full sample) were also included in the model but their coefficients are not shown.

| | $\beta$ | SE($\beta$) | z value | Pr(>\|z\|) |
| --- | --- | --- | --- | --- |
| (Intercept) | 15.4059 | 1.6104 | 9.57 | $< 2 \times 10^{-16}$ |
| N.ROH2 | 0.0185 | 0.0155 | 1.19 | 0.2332 |
| N.ROH1 | -0.0351 | 0.0093 | -3.78 | 0.0002 |
| HET | -42.3276 | 4.4278 | -9.56 | $< 2 \times 10^{-16}$ |
| Consanguinity score = 1 | 0.3102 | 0.1966 | 1.58 | 0.1146 |
| Consanguinity score = 2 | 0.5183 | 0.1435 | 3.61 | 0.0003 |

**Table S2:**
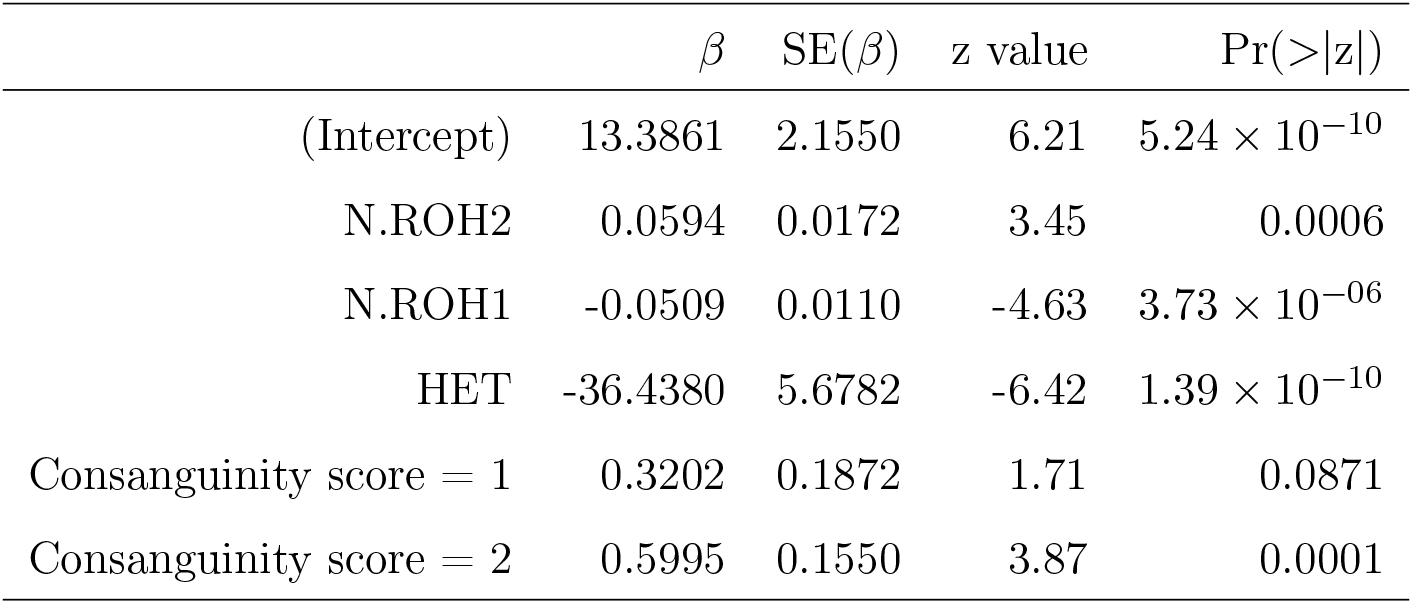
Poisson regression model fitted for NROH10 as response (population - weighted). Ten genetic PCs (calculated for the full sample) were also included in the model but their coefficients are not shown.

**Table S3:** Poisson regression model fitted for NROH10 in South Asia (population - unweighted). Ten genetic PCs (calculated in South Asians only) were also included in the model but their coefficients are not shown.

| | $\beta$ | $SE(\beta)$ | z value | $Pr(>\vert z\vert )$ |
| --- | --- | --- | --- | --- |
| (Intercept) | 27.1465 | 3.2748 | 8.29 | $< 2 \times 10^{-16}$ |
| N.ROH2 | 0.0277 | 0.0381 | 0.73 | 0.4668 |
| N.ROH1 | -0.0343 | 0.0208 | -1.65 | 0.0983 |
| HET | -75.5652 | 9.0313 | -8.37 | $< 2 \times 10^{-16}$ |
| Consanguinity score = 1 | 0.1002 | 0.3389 | 0.30 | 0.7675 |
| Consanguinity score = 2 | 0.4922 | 0.2415 | 2.04 | 0.0415 |

**Table S4:**
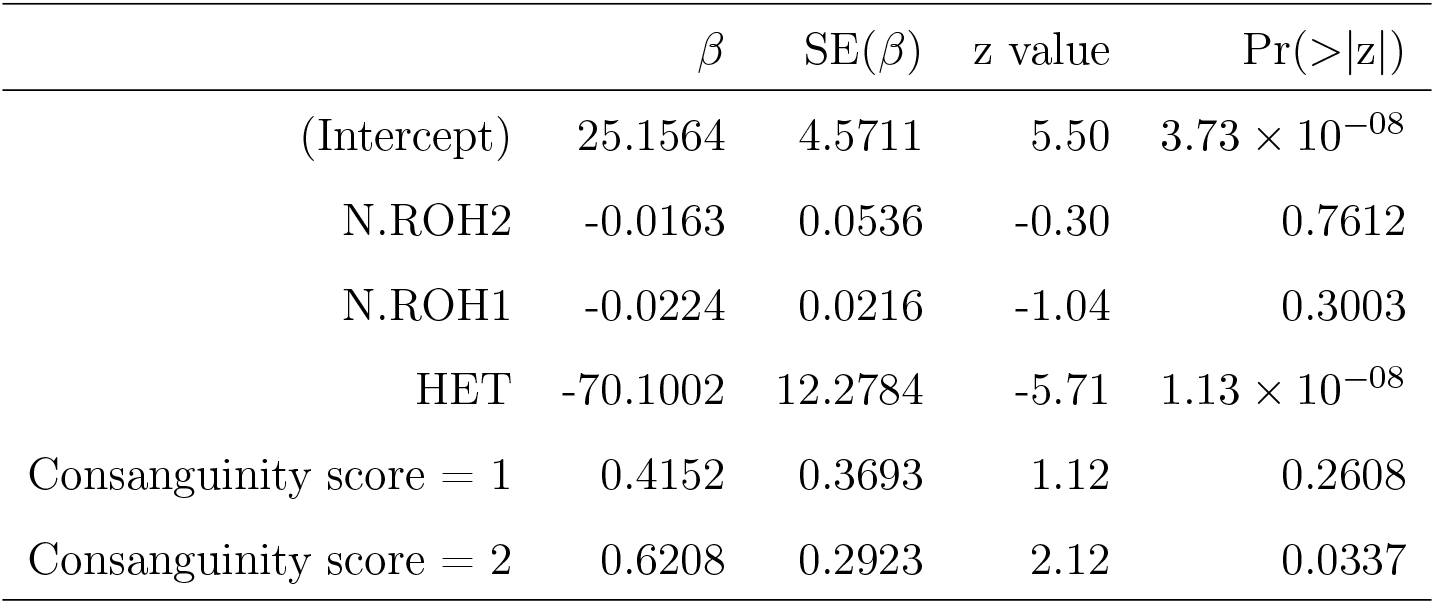
Poisson regression model fitted for NROH10 in South Asia (population - unweighted). Ten genetic PCs (calculated in South Asians only) were also included in the model but their coefficients are not shown.

**Table S5:** Effect of consanguinity on NROH10 in South Asia. Poisson regression model (individual-level) with NROH10 used as response. Coefficients for consanguinity score is shown relative to the score of 0 (consanguinity prohibited). Coefficients of 1 and 2 represent populations which permit and prefer cousin unions, respectively. Ten genetic PCs (calculated in South Asians only) were also included in the model but their coefficients are not shown.

| | $\beta$ | $SE(\beta)$ | z value | $Pr(>\vert z\vert )$ |
| --- | --- | --- | --- | --- |
| (Intercept) | 19.8837 | 0.5459 | 36.43 | $< 2 \times 10^{-16}$ |
| N.ROH1 | -0.0207 | 0.0038 | -5.51 | $3.6 \times 10^{-08}$ |
| N.ROH2 | -0.0170 | 0.0058 | -2.91 | 0.0036 |
| HET | -56.3922 | 1.4404 | -39.15 | $< 2 \times 10^{-16}$ |
| Consanguinity score = 1 | 0.3082 | 0.1212 | 2.54 | 0.0110 |
| Consanguinity score = 2 | 0.5992 | 0.0977 | 6.13 | $8.68 \times 10^{-10}$ |

**Table S6:** Genetic differences between pairs of populations preferring and prohibiting consanguinity (outliers removed, N = 68 pairs)

|  | Variable | Statistic | Pvalue |
| --- | --- | --- | --- |
| 1 | NROH10 | 1155.50 | 0.04 |
| 2 | NROH5 | 1155.50 | 0.75 |
| 3 | NROH2 | 1191.00 | 0.91 |
| 4 | NROH1 | 1091.00 | 0.62 |
| 5 | HET | 969.00 | 0.21 |

